# The causal role of three frontal cortical areas in grasping

**DOI:** 10.1101/2020.06.25.170126

**Authors:** I Caprara, P Janssen

## Abstract

Efficient object grasping requires the continuous control of arm and hand movements based on visual information. Previous studies have identified a network of parietal and frontal areas that is crucial for the visual control of prehension movements. Electrical microstimulation of 3D shape-selective clusters in AIP during fMRI activates areas F5a and 45B, suggesting that these frontal areas may represent important downstream areas for object processing during grasping, but the role of area F5a and 45B in grasping is unknown. To assess their causal role in the frontal grasping network, we reversibly inactivated 45B, F5a and F5p during visually-guided grasping in macaque monkeys. First, we recorded single neuron activity in 45B, F5a and F5p to identify sites with object responses during grasping. Then, we injected muscimol or saline to measure the grasping deficit induced by the temporary disruption of each of these three nodes in the grasping network. The inactivation of all three areas resulted in a significant increase in the grasping time in both animals, with the strongest effect observed in area F5p. These results not only confirm a clear involvement of F5p, but also indicate causal contributions of area F5a and 45B in visually-guided object grasping.

## 1. Introduction

To grasp an object, the brain needs to have a fine representation of its shape in order to adjust the hand efficiently before contact. For this purpose, visual information needs to be processed in several aspects. The object has to be located in space, and analyzed in terms of geometry and orientation through the dorsal visual stream (Janssen and Scherberger, 2015; Theys et al., 2015). Extensive research has investigated which areas along the dorsal stream are involved in visual object processing for motor preparation. Area AIP, in the rostral lateral bank of the intraparietal sulcus (IPS), supports goal-directed hand movements (Taira et al., 1990; Murata et al., 2000; Baumann et al., 2009). AIP neurons are selective for specific grasp types and preserve the same object preference during fixation. Gallese et al. (1994) observed a clear grasping deficit after reversible inactivation of AIP, while the inactivation of LIP only led to misreaching. In frontal cortex, Fogassi et al. (2001) used the same approach to investigate the causal role of area F5 in the grasping network by injecting muscimol in the posterior bank of the arcuate sulcus – F5p – and in the convexity – F5c. The reversible inactivation of F5p led to a significant increase in the grasping time and an impairment of the hand preshaping, while much milder muscimol effects were observed when inactivating area F5c. However, based on architectonic criteria and cortical connectivity, Belmalih et al. (2009) and Gerbella et al. (2011) described a third and more anterior subsector, F5a. Neurophysiological studies have demonstrated that F5a neurons are selective for three-dimensional (3D) stimuli (higher order disparity), and are mainly active during visually guided - but not during memory-guided grasping, i.e. visual-dominant neurons (Theys et al., 2012a; Theys et al., 2012b). However, direct causal evidence for a role of F5a in visually-guided object grasping is currently lacking. Area 45B, a neighboring area located in the anterior bank of the arcuate sulcus, is connected to the posterior subsector of AIP, as demonstrated with electrical microstimulation during fMRI (Premereur et al., 2015; Caprara et al., 2018), which also suggests a role in the visuomotor grasping network. Although Gerbella et al. (2010) initially hypothesized an involvement in the oculomotor system similar to the Frontal Eye Fields (FEF), the ability of 45B neurons to encode images of objects has been extensively documented (Caprara et al., 2018). It remains unclear whether neurons in 45B are part of a more extended grasping network, together with the different subsectors of F5.

To our knowledge, no study has systematically tested the causal role of area F5a and area 45B in object grasping. We performed reversible inactivations in area F5 (anterior and posterior subdivisions) and in area 45B during visually-guided object grasping. We hypothesized thatif area 45B and F5a are involved in object grasping, we would observe a behavioral deficit as previously described after F5p inactivation.

## 2. Materials and Methods

### 2.1 Subjects

Two adult male rhesus monkeys (D, 7 kg; Y 14 kg) were used as subjects for this study. All experimental procedures, surgical techniques, and veterinary care were performed in accordance with the NIH Guide for Care and Use of Laboratory Animals and in accordance with the European Communities Council Directive 2010/63/EU, and were approved by the local ethical committee on animal experiments of the KU Leuven.

An MRI-compatible head fixation post and recording chamber (Crist Instruments, Hagerstown, MD), were implanted under propofol anesthesia using dental acrylic and ceramic screws above the right and left arcuate sulcus in Monkey D, and in Monkey Y, respectively.

### 2.2 Apparatus

During the experiments, monkeys were seated upright in a chair with the head fixed. The animals were previously trained not to move the ipsilateral arm during the whole duration of the session. An industrial robot (Universal Robots) presented one of two different objects (a large and a small sphere – 3 cm and 1 cm diameter respectively) in front of the monkey (Universal Robots), in blocks of 10 trials, at a reachable position (36 cm, at chest level – ~ 20 cm reaching distance measured from the center of the hand rest position to the center of the objects – Figure 1A). Both objects drove a pad-to-side/scissor combination of grip types (Napier, 1993; Macfarlane and Graziano, 2009).

**Figure 1.**
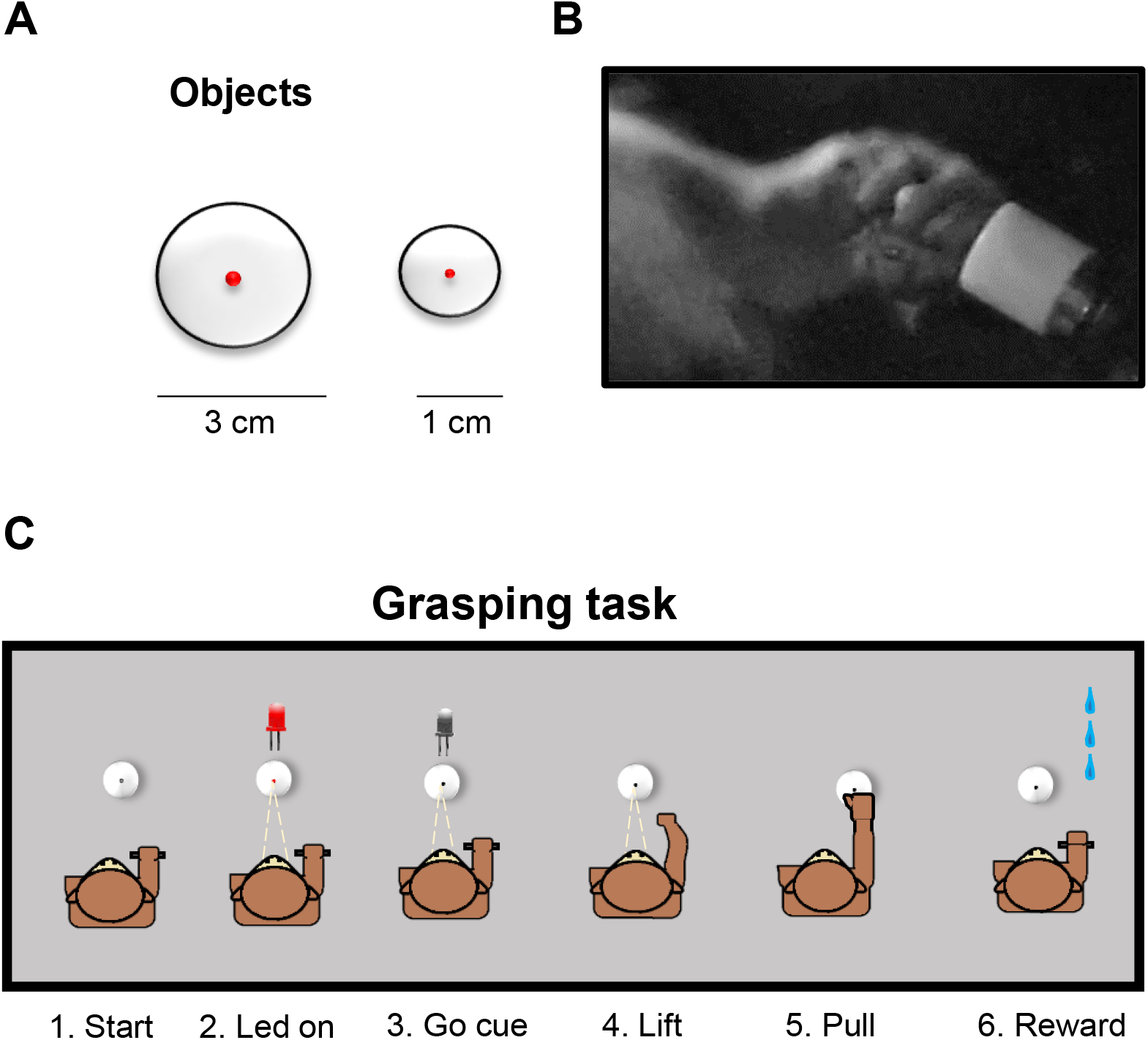
Stimuli, grip type and behavioral task. A. Objects. Large and Small sphere (3 and 1 cm of diameter, respectively). B. Grip type. Monkey D combination of scissor/and pad to side grip, mainly using third, fourth and fifth finger; same grip was used by Monkey Y (not shown). C. Behavioral task, performed in a semi-dark room. The trial starts with the monkey holding its hand on the resting position (1). A red LED in the center of the objects becomes bright (2), and the monkey holds fixation for a variable time window (850-2000 ms). Then, the LED dims, serving as a Go cue (3) to lift the hand (4), reach and pull (5) the object. If the trial is performed correctly, the animal receives few drops of reward.

Fiber-optic cables detected the resting position of the hand, the start of the reach to grasp movement, and the pulling of the object. The start of the hand movement was detected as soon as the palm of the hand was 0.3 cm above the resting plane, whereas pulling of the object was detected when the object was pulled for 0.5 cm in the horizontal axis. We continuously monitored the position of the left eye with an infrared-based camera system (Eye Link II, SR Research) sampling pupil position at 250 Hz.

### 2.3 Behavioral Task

Each monkey performed a visually-guided grasping (VGG) task in a semi-dark room (Figure 1C). The trial started when the monkey placed the contralateral hand in the resting position. During this time, the robot picked an object from a box and presented it at a reachable distance (36 cm for Monkey D; 28 cm for Monkey Y). A red LED inserted in the middle of the object was illuminated, which the monkey had to fixate, keeping the gaze inside a ±3.5 degree fixation window. After a variable amount of time (850-2000 ms), the red LED dimmed, serving as ‘Go cue’ for the monkey to lift its hand from the resting position (within 2500 ms from the ‘Go cue’), reach and pull the object for 300 ms (holding time). Because the monkeys performed the experiments in a dimly lit room, the object was clearly visible in all epochs of the task. If all of these stages were accomplished correctly, the monkey would obtain few drops of reward.

### 2.4 Reversible inactivation procedure

In each of the three areas (45B, F5a and F5p), we first performed extensive recordings of singleunit activity during visually guided grasping of four different objects (a large sphere, a small sphere, a large plate and a small plate) presented by the robot. In these recordings, the animal initiated the trial in the dark with the hand in the resting position, after which a red LED appeared on the object as a fixation point. After 500 ms of fixation, a white light illuminated the object from within. Then, after a variable delay of 350 ms to 1500 ms, the red LED dimmed, which was the go-cue for the monkey to grasp and pull the object. After having identified the recording sites with the strongest object responses, we estimated the optimal depth for the muscimol injections. To visualize the spread of the drug prior to the inactivation sessions, we first injected 4 μl of a contrast agent (Dotarem, 2% solution) with a 10 μl Hamilton syringe inserted into a stainless steel guiding tube placed in the grid (Figure 2), and obtained anatomical Magnetic Resonance Images at 0.6 mm resolution.

**Figure 2.**
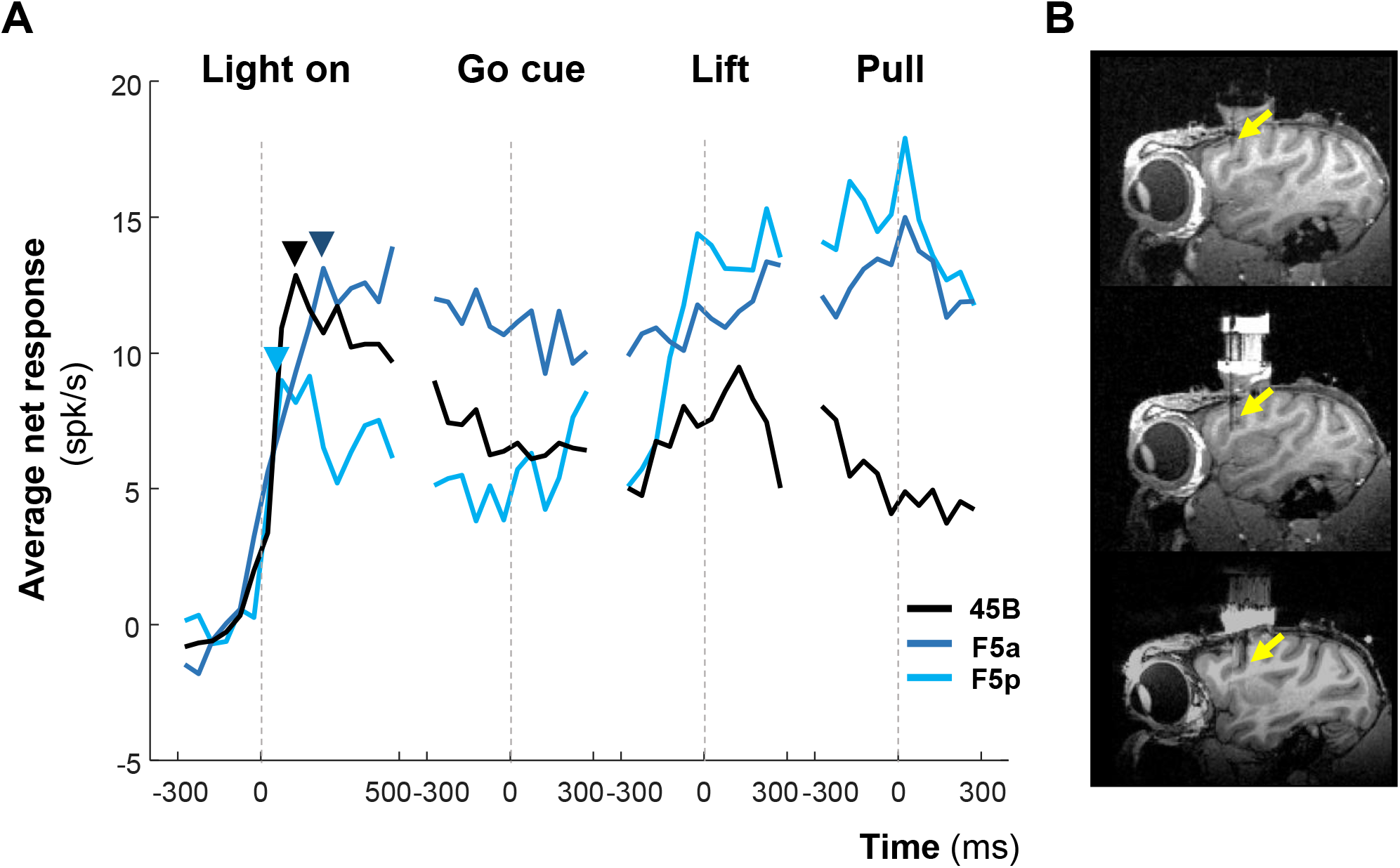
Average net grasping activity recorded in area 45B (N = 114), F5a (N = 89) and F5p (N = 77) during a visually guided grasping task. A. Average response Data are aligned to Light on (−500/+300 ms); Go cue (−300/+300 ms); Lift (-300/+300 ms); Pull (−300/+300 ms). Bin width is 50 ms. B. Sagittal view of the Monkey D best recording position in area 45B, F5a and F5p, respectively. Yellow arrows represent the electrode tip. Comparable positions were also targeted in Monkey Y (data not shown).

To reduce cortical damage induced by the injection, we lowered the tip of the needle until the transition between the white and the grey matter (in the case of 45B) or at the level of the sulcus (F5a). One week after the Dotarem control, we started the reversible inactivation experiments. To measure the normal behavior of the animal, each session started with at least 30 minutes of behavioral data collection during the execution of the task (‘Pre’). Then, after the Pre period, we injected 4μl of GABA-A agonist muscimol (Sigma, 10 mg/ml – same procedure as for the Dotarem control – ‘Post’). Behavioral data were collected immediately after the injection until the monkey stopped working (minimum 2 hours after the injection).

To avoid pressure damage, the infusion in each site was slowly delivered (1 μl/min). We performed at least one control session in each area (except for F5a in monkey Y due to a health problem), injecting the exact same volume of saline (4 μl) in the same grid position. Saline sessions were carried out after a sequence of three inactivation sessions. Each session – muscimol or saline – was followed by at least 48h where no injection was performed.

Finally, to obtain reference data with which we could compare the effect of our inactivations, we used the same protocol to inactivate primary motor cortex (M1), whose impairment has been extensively described (Brochier et al., 1999; Fogassi et al., 2001; Stepniewska et al., 2014). For this control injection, we used the most posterior grid position in the same recording chamber of Monkey D. The movements of the hand were video recorded in each experimental session.

### 2.5 Data Analysis

All data analyses were performed in Matlab (Mathworks).

In order to quantify the performance of the monkey, we calculated the time between the Go cue and the pull of the object (Total Grasping time – TT). For a more detailed analysis of the behavior, we divided the trial into two epochs: the reaction time (RT – from Go cue to Lift of the hand) and grasping time (GT) – from Lift of the hand to Pull. We calculated the percentage change in RT and GT for each area (Variation index – *vi*).

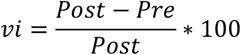

To investigate the reproducibility of the inactivation effect over sessions, we plotted the behavioral performance (RT, GT, TT) before and after muscimol in the inactivation sessions, per area. To test the significance of the muscimol effect, we compared the performance from 30 min after the injection until the end of the session with that before the injection (unpaired t-test, p<0.01).

Then, to exclude aspecific effects caused by the injection, we compared the average TT across muscimol sessions with that of the saline session (unpaired t-test, p<0.01). Finally, in order to have a complete description of the animal’s behavior, we also calculated the percentage error trials (errors occurring after the Go cue) after muscimol injection and compared them to those of the Pre periods in each session. We also tested whether the muscimol injections had any effect on the eye position by calculating the variance of the eye position (both x and y position) before and after muscimol injection, separately for the three areas.

## Results

### Single-cell responses during grasping in area 45B, F5a and F5p

Figure 2 shows the average neural activity recorded in the three targeted areas during visually-guided grasping of the four objects (the object selectivity in the three areas is the topic of a separate study). In each area, we observed distinct responses during grasping. Area 45B (N = 114) showed a fast visual response to Light onset in the object, reaching a maximum at 100-150 ms (bin width of 50 ms). Then, after a small decrease in activity before the go-cue, we observed a weak response around the lift of the hand (p = 0.08, pre vs post-lift epoch), and reduced activity (compared to the object fixation epoch, p = 0.01) around the pull of the object at the end of the trial. F5a neurons (N = 89) had a slower onset response (peak at 200-250 ms after Light onset), the activity remained high throughout the whole duration of the trial and peaked at the pull of the object. Surprisingly, the population activity in area F5p (N = 77) showed a fast visual response to Light onset (maximum at 50-100 ms after Light onset), followed by a relatively large drop in activity during the delay period before the go cue, and a strong increase in activity before and during the hand movement (p = 3.07 x 10^−6^, pre vs post-lift epoch), with a maximum at the pull of the object. Thus, all three areas of interest showed robust responses during visually-guided object grasping with distinct temporal dynamics. Area 45B neurons exhibit strong visual responses (in line with our previous study, Caprara et al. 2018), but do not modulate their activity much around the lift of the hand, whereas F5p neurons show the strongest response modulation before and during the hand movement.

### Effect of reversible inactivation of area 45B, F5a and F5p

To obtain a reference value for the behavioral deficit in our task after muscimol injection, we injected 4 μl of muscimol in the arm/hand region of M1 (in the anterior bank of the central sulcus, Figure 3A), whose inactivation effect has been extensively described in the literature (Brochier et al., 1999; Fogassi et al., 2001; Stepniewska et al., 2014). TT, RT and GT increased significantly after muscimol injection (*t-test*, p<0.01), with an increase of 31% in TT compared to pre-muscimol testing (*z-test*, p < 0.001 – Figure 3B). Note however that the animal was capable of performing the task (584 correct trials compared to 159 error trials in 120 min of testing). To further assess the level of impairment after muscimol, we gave the monkey the opportunity to grasp a small nut – smaller than the 1 cm sphere presented by the robot – from the experimenter hand at the end of the M1 inactivation session. We observed a clear paresis of the fingers of the contralateral hand, resulting in a strong grasping deficit, which prevented the monkey from picking up the fruit (Figure 3C). Additionally, when coming back to the resting position, the hand was placed in an unnatural way, often characterized by an odd wrist orientation.

**Figure 3.**
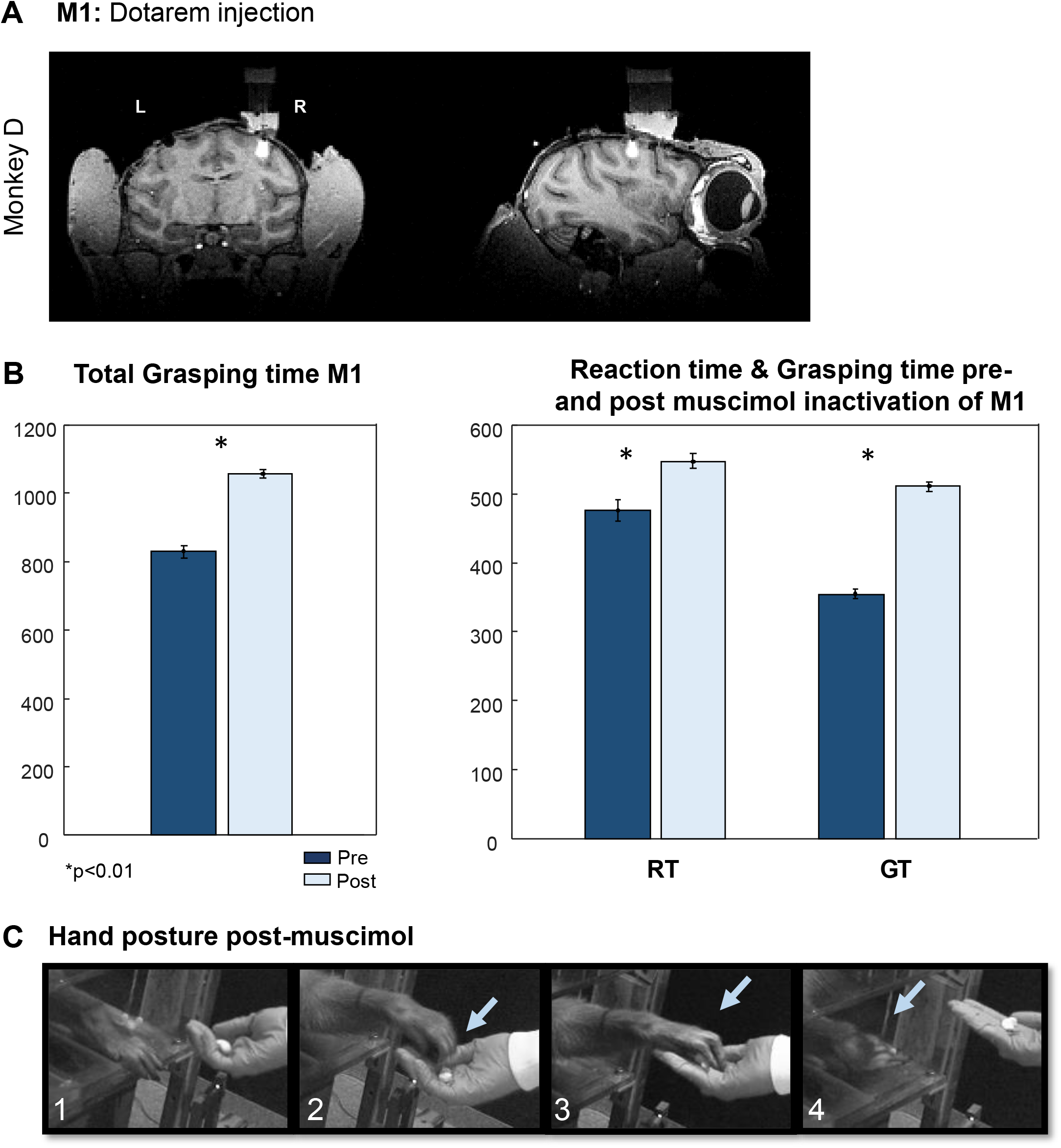
Inactivation of M1. A Dotarem injection. 4μl of 2% Dotarem injected in the hand-related subregion of M1. B. Total grasping time (TT), Reaction time (RT) and Grasping time (GT) across sessions. Left panel: TT, Pre and Post muscimol injection (dark and light blue, respectively); Right panel: RT, and GT. Asterisks indicate significance (p<0.01). C. Grasping sequence showing the hand posture after muscimol injection. The photographic sequence has been taken at the end of the session, while the Monkey D attempted to grasp a nut from the observer’s hand. Light blue arrows represent the unnatural posture of the hand: panels 2-3 show lack of strength in the fingers; panel 4 shows the unnatural posture of the hand at the end of the attempt.

We then verified the locations of our injections in the three frontal areas of interest by means of dotarem injections in the scanner. Each injection targeted the recording sites which gave strong task-related responses during grasping. The 45B injection was centered on the anterior bank of the arcuate sulcus (Figure 4B), covering a volume of approximately 4 (anterior-posterior) by 4 (medio-lateral) by 6 (dorso-ventral) mm. The F5a injection was centered on the posterior bank of the arcuate sulcus, 1 mm more anterior and 3 mm deeper (more ventral) than the 45B injection, with a similar volume of cortex covered. The F5p injection was largely confined to the ascending part of the posterior bank of the arcuate sulcus, 4 mm more posterior compared to the 45B injection. The overlap between the 45B and the F5a injection, and between the 45B and the F5p injection was minimal (Figure 4C), while we did not observe any overlap between the F5a and the F5p injections. We obtained highly similar results in monkey Y (data not shown). Thus, despite the close proximity of the three frontal areas to each other, we achieved highly targeted inactivations with minimal overlap.

**Figure 4.**
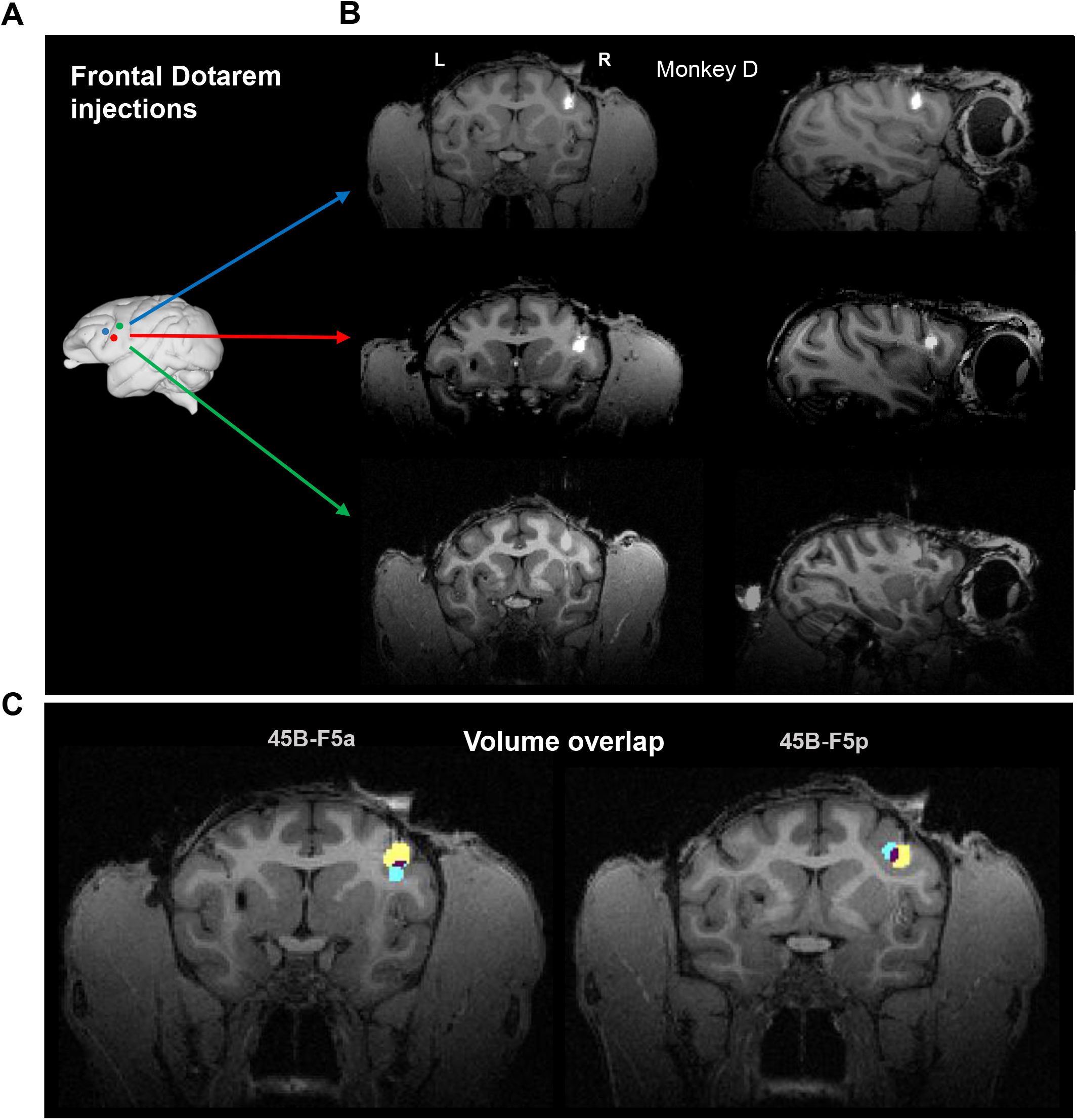
Dotarem injections. A. Anatomical location of areas of interest. Brain image was edited from the ‘Scalable Brain Atlas’ (http://link.springer.com/content/pdf/10.1007/s12021-014-9258-x; Calabrese et al 2015). B. 4μl of Dotarem (2%) injected respectively in area 45B, F5a and F5p of Monkey D. C. Coronal section of arcuate sulcus in Monkey D, showing the overlap (represented in purple) between Dotarem injections of area 45B (represented in yellow) with F5a (represented in cyan), and F5p (represented in cyan), respectively. Area F5a and F5p positions of injection, instead did not present any overlap (not shown).

In a first analysis, we combined the data obtained with the large and the small sphere (Figure 5). Reversible inactivation of area F5p induced a significant increase in TT in both monkeys (*t-test*, p = 9.38 x 10^−14^ and 9.49 x 10^−63^, for Monkey D and Monkey Y, respectively – Figure 5), in line with previous studies (Fogassi et al., 2001). However, we also observed a significant effect on TT following the inactivation of area 45B and area F5a (*t-test*, p = 6.71 x 10^13^ and 1.27 x 10-^06^ for 45B and F5a in Monkey D; p = 8.54 x 10^−29^ and 3.66 x 10^−32^ for 45B and F5a in Monkey Y). The effect of muscimol on TT differed significantly between the three areas (twoway ANOVA with factors *area* and *muscimol*) in Monkey D (p = 9.90 x 10^−3^) but not in monkey Y (p = 0.42). However, the magnitude of the muscimol effect on TT was relatively comparable across the three areas (20% increase for 45B, 19% increase for F5a and 22% increase in F5p, averaged across the two monkeys).

**Figure 5.**
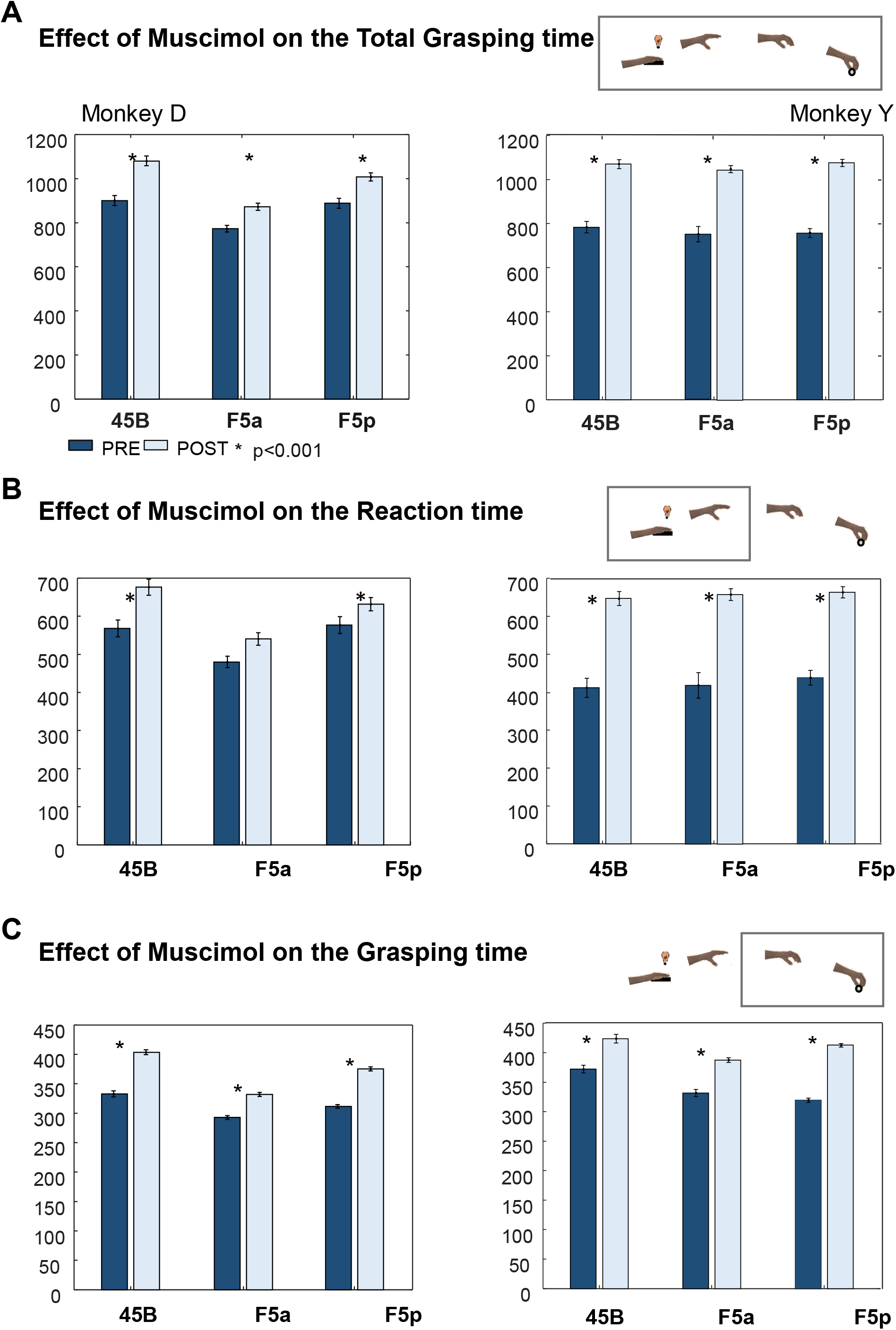
TT, RT and GT for the three frontal areas averaged across sessions. The bar plots indicate the Pre period (before inactivation) in dark blue, while the Post inactivation (from 30’ after muscimol injection until end of the session) is represented in light blue, for Monkey D (left panels) and Monkey Y (right panels). Asterisk indicates significance with p<0.001.

Next, we investigated which epoch of the trial was the most affected by reversible inactivation by subdividing the TT into an RT (Reaction time; the time between the go-cue and the lift of the hand) and a GT (Grasping time; the time between the lift of the hand and the pull of the object). In all three areas, we measured a significant increase in both RT and GT (*t-test*, all p-values <0.001), with the exception of RT after F5a inactivation (Figure 5B).

To quantify the effect size 30 minutes after muscimol injection, we calculated the percentage change (See Methods 3.5) in RT and GT after reversible inactivation of each of the three targeted areas (Table 1). The highest percent change in the GT after muscimol injection was observed in F5p (Monkey D: 19%; Monkey Y: 22%), whereas weaker effects were observed in area 45B (Monkey D: 15%; Monkey Y: 11%) and F5a (Monkey D: 14%; Monkey Y: 16%). Overall, the GT behavioral effects we observed in the three frontal areas were robust but significantly smaller than those observed after M1 inactivation: a two-way ANOVA with factors *muscimol* (pre-post) and *area* showed significant interaction effects in Monkey D (p = 3.70 x 10^−16^ for the 45B-M1 comparison, p = 1.61 x 10^−12^ for the F5p-M1 comparison and p = 2.26 x 10^−28^ for the F5a-M1 comparison).

**Table 1.** Variation index: performance percent change after muscimol in Reaction time (RT) and Grasping time (GT). See methods for detailed explanation.

|  | 45B |  | F5a |  | F5p |  |
| --- | --- | --- | --- | --- | --- | --- |
|  | Monkey D | Monkey Y | Monkey D | Monkey Y | Monkey D | Monkey Y |
| RT | 14% | 36% | 6% | 37% | 11% | 34% |
| GT | 15% | 11% | 14% | 16% | 19% | 22% |

In order to verify whether the deficits we observed were caused by the temporary inactivation of the areas rather than by aspecific factors (e.g. fatigue or a decrease in motivation because of the injection procedure), we performed one saline session in each area in Monkey D., and one saline session in area 45B and F5p in monkey Y. As expected, similar to previous studies (Demer and Robinson, 1982; Fogassi et al., 2001), muscimol induced significantly longer GTs than saline in each area (*t-test*, p<0.001 – Figure 6). We also performed a single (4 μl) muscimol injection in area 46 in prefrontal cortex in monkey D (3 mm more anterior than the 45B injection – Figure 7). In contrast to the other frontal areas, this injection in area 46 mainly caused an increase in RT (*t-test*, p = 4.32 x 10^−6^), and a smaller effect on the GT (*t-test*, p = 3.60 x 10^−3^), which was significantly smaller than the effect in 45B (*t-test*, p = 0.041).

**Figure 6.**
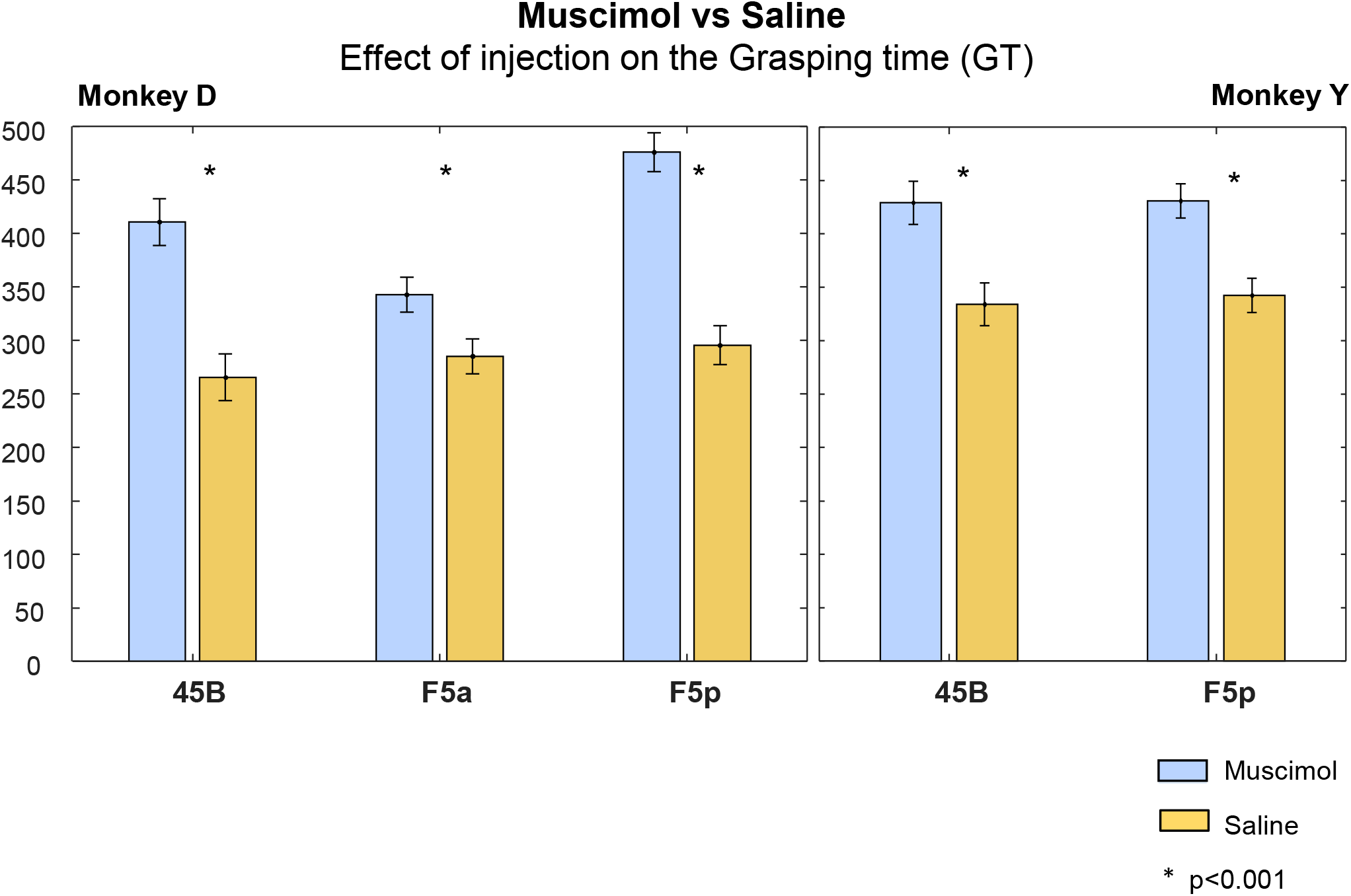
Effect of muscimol vs Effect of Saline. Monkey D tested for area 45B, F5a and F5p (left panel); Monkey Y tested for 45B and F5p (right panel). Light blue corresponds to Muscimol (Session 1); yellow corresponds to the effect of Saline. Asterisk indicates significance with p<0.001.

**Figure 7.**
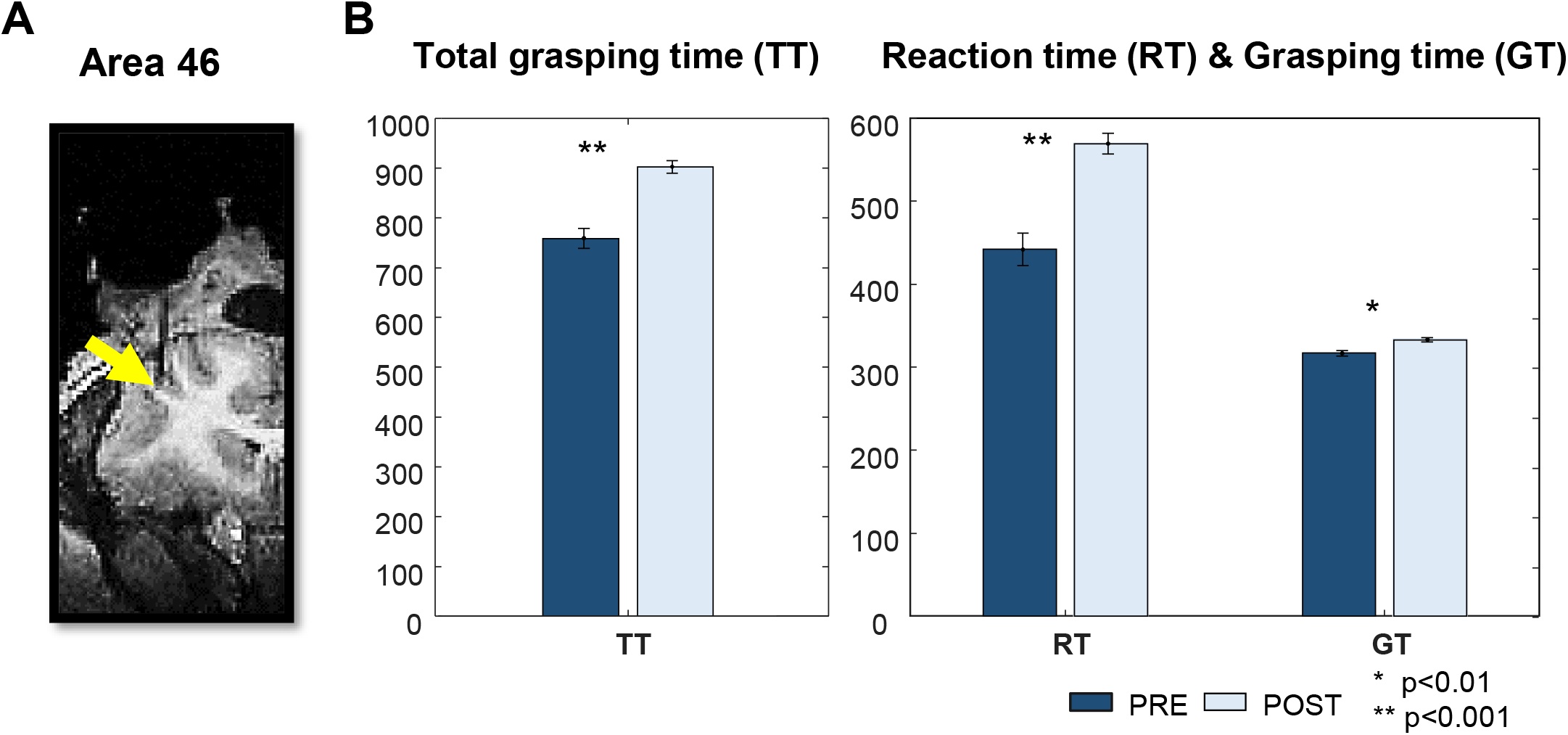
Inactivation of Area 46. A. Injection position. Black trace represents an electrode lowered in the position of muscimol injection in Area 46. B. Grasping Time (GT), Reaction Time (RT) and Total Grasping Time (TT) across session – Pre and Post muscimol injection (dark and light blue, respectively). One asterisk indicate significance with p<0.01; two asterisks indicate p<0.001.

The inactivation of these three areas in the arcuate sulcus did not lead to obvious visible impairments during grasping. However, the reversible inactivation of area F5p induced a mild change of the grip type (Figure 8), from a combination of pad-to-side/scissor grip to a more unnatural posture of the fourth and fifth finger. Instead, the inactivation of area 45B and of F5a did not lead to visible deficits in the hand preshaping. Despite this, in all three cases, the monkey was capable of reaching and grasping the object quite efficiently (percentage correct trials in Monkey D – 45B: 44%, F5p: 54%, F5a: 44%; Monkey Y – 45B: 34%, F5p: 53%, F5a: 47%).

**Figure 8.**
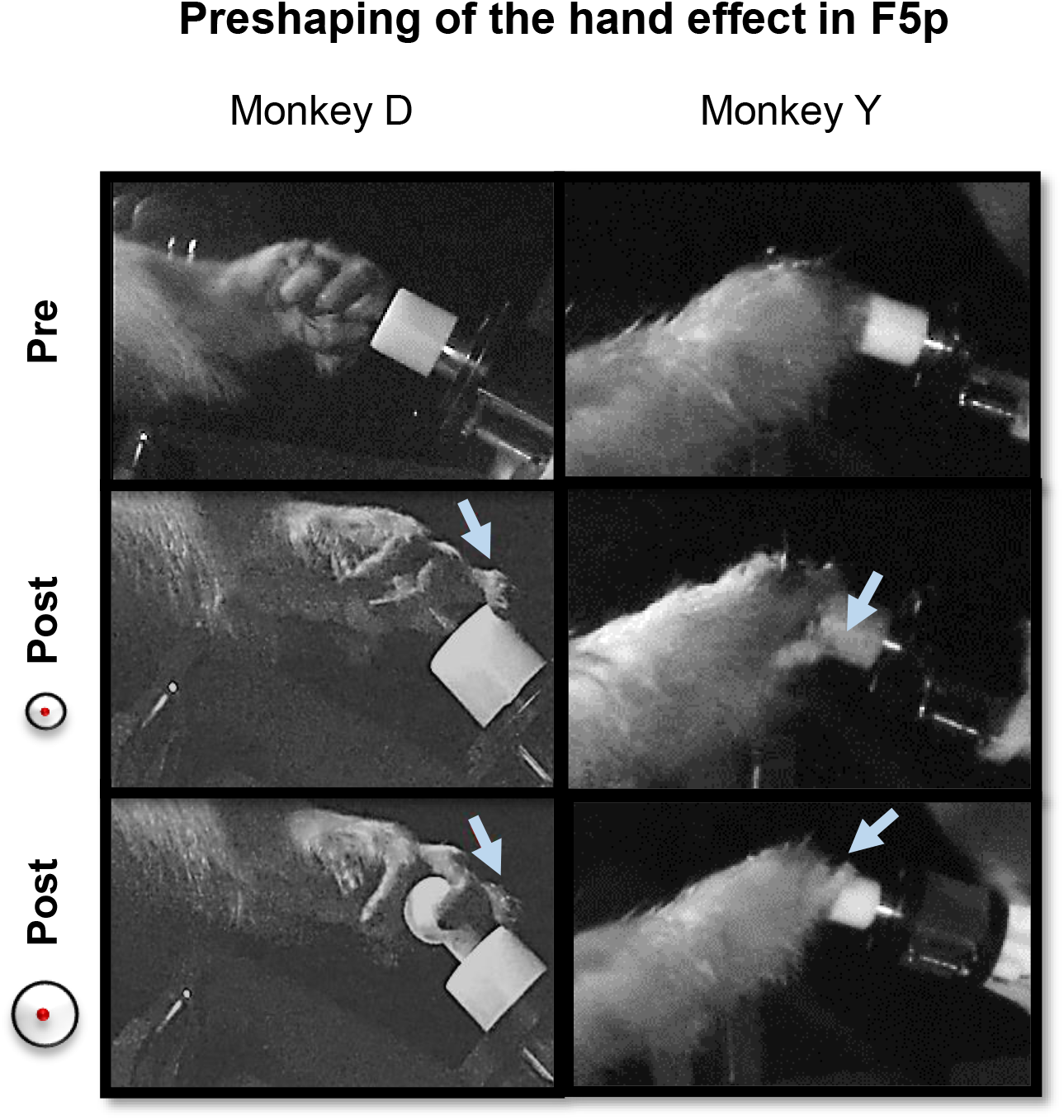
Grip after muscimol injection in area F5p. Odd posture of the fourth and fifth fingers in Monkey D, and of the fifth finger in Monkey Y, during the grip of 1 and 3 cm spheres diameter (upper and lower panel, respectively), accompanied by a less powerful and sloppier grasping.

We also investigated the behavioral effect of muscimol across sessions (Figure 9). Monkey Y showed highly consistent effects of muscimol on TT (8 out of 9 sessions with a significant effect), while monkey D showed effects that were more variable over sessions (5 out of 9 session with a significant effect on TT). Although some of this variability in Monkey D could be related to speed-accuracy tradeoffs (e.g. session 2 in F5a with long RTs and no effect on GT), other sessions (e.g. session 2 and 3 in F5p) did not exhibit any effect of muscimol injection. Note that this intersession variability in F5p could not be caused by short intervals between injections, since we left 20 days between the first and the second session and 3 days between the second and the third session, and the effect of muscimol was completely reversed within 48h.

**Figure 9.**
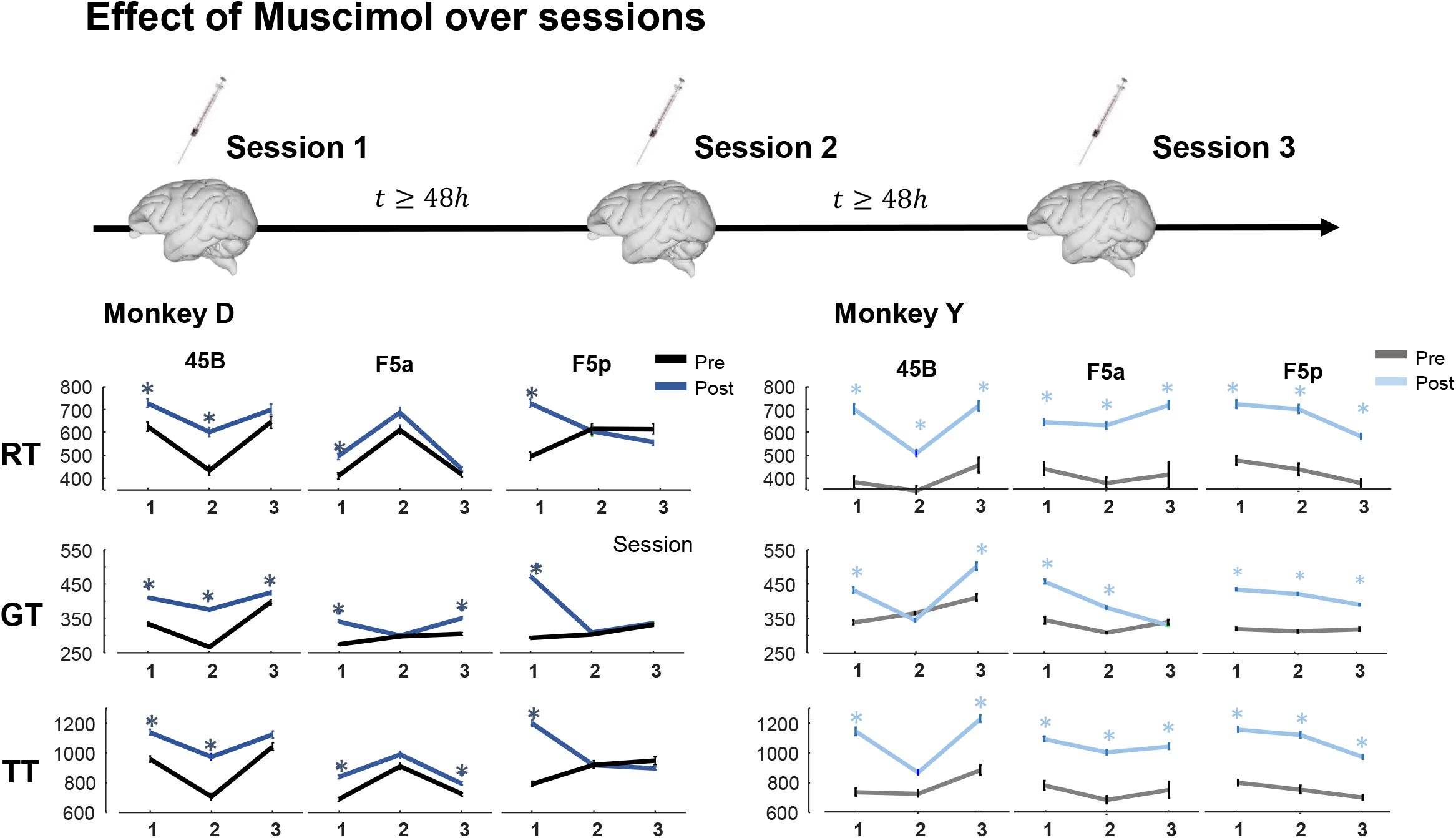
Effect of muscimol over three sessions on RT, GT and TT. Monkey D (left panel): dark blue represent the period after inactivation (Post), and the black lines represent the period before (Pre). Monkey Y (right panel): light blue represents the Post inactivation, while grey represents Pre inactivation. Sessions were spaced in time of minimum 48h to guarantee that muscimol effect was completely reversed. Asterisk represents a p<0.01.

We also examined the effect of muscimol inactivation on the two objects that had to be grasped (Figure 10). Muscimol inactivation of all three areas significantly increased the GTs for both the large and the small object in both animals. Then, we tested whether the effect of inactivation differed between the objects by comparing the changes in GT (relative to the pre-muscimol trials) between the two objects. The injection of muscimol in 45B and F5p (Monkey D) and in 45B and F5a (Monkey Y) prolonged the GTs much more for the small object compared to the large object (*t-test*, 1.48 x 10^−9^ and 0.010 respectively in Monkey D and 1.48 x 10^−9^ and 6.04 x 10^−26^ respectively in Monkey Y).

**Figure 10.**
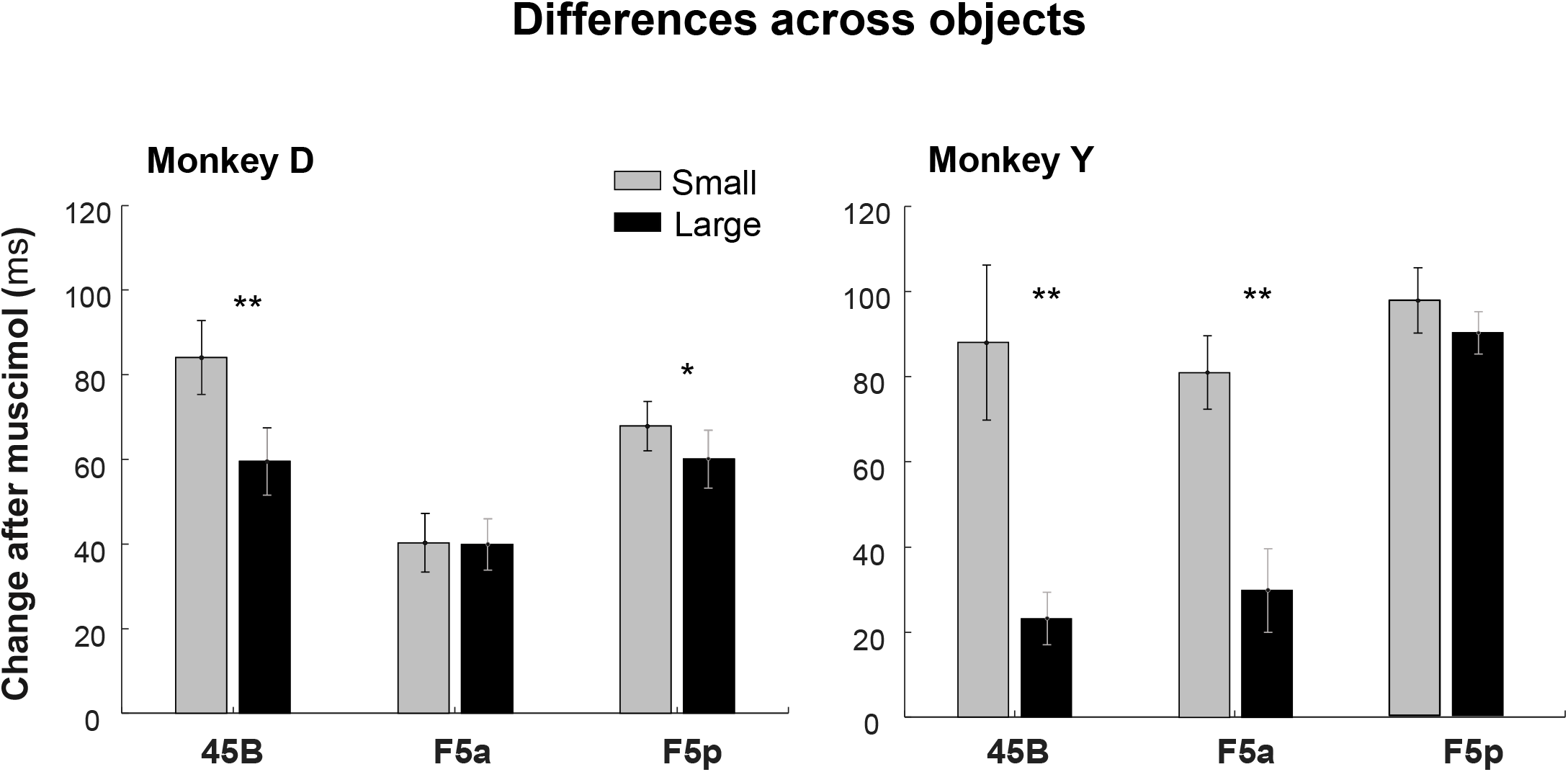
Average change in GT across session for the two objects. Grey bars correspond to the small sphere (1 cm); Black bars correspond to the large sphere (3 cm). One asterisk indicates p<0.05; two asterisks, p<0.001.

To test whether the accuracy of performance changed after the muscimol injection, we calculated the number of errors made by the monkey after the Go cue event until the end of the grasping trial (the pull of the object). Reversible inactivation of all three areas induced small but significant increases in the number of errors (Table 2).

**Table 2.** Accuracy of performance, before (Pre) and after (Post) muscimol injections. Percentages represent the number of errors made by the monkeys in from the Go cue event until the end of the grasping trial (the pull of the object). Grey columns represent significant increase in the number of errors after muscimol injection (z-test; p<0.01).

|  | 45B |  | F5a |  | F5p |  |
| --- | --- | --- | --- | --- | --- | --- |
|  | Monkey<br>D | Monkey<br>Y | Monkey<br>D | Monkey<br>Y | Monkey<br>D | Monkey<br>Y |
| Pre | 3.9% | 2.3% | 5.2% | 0.5% | 6.5% | 0.3% |
| Post | 8.2% | 5.7% | 4.0% | 1.6% | 3.8% | 1.1% |

In view of the proposed role of area 45B in oculomotor control (Gerbella et al., 2010) we investigated changes in eye fixation following the muscimol injections, since a longer GT could result from difficulties in object fixation. To this end, we calculated the trial-to-trial variability of the eye position in the fixation epoch (i.e. 200-500 ms after the Led onset) before and after muscimol. In Monkey D, the eye position variance increased significantly after muscimol injection in area F5a and in area F5p (both x and y position in F5a and x position in F5p; *F-test*, Table 3), while in area 45B it decreased (only the y position). Monkey Y, instead, only showed a significant decrease in eye position variance after F5a and F5p inactivations (p = 7.03 x 10^−6^ Y-coordinate and p = 7.60 x 10^−4^, X-coordinate, respectively) but not in 45B (Table 3). Thus, the analysis of the eye position variance suggests that the deficit we observed in 45B was most likely not caused by an oculomotor effect. In addition, monkey Y showed a clear grasping deficit after 45B inactivation in the absence of a change in eye position variability. Finally, the proportion of error trials caused by a break of fixation during the trial did not increase significantly after muscimol injection in the large majority of the sessions (data not shown).

**Table 3.**
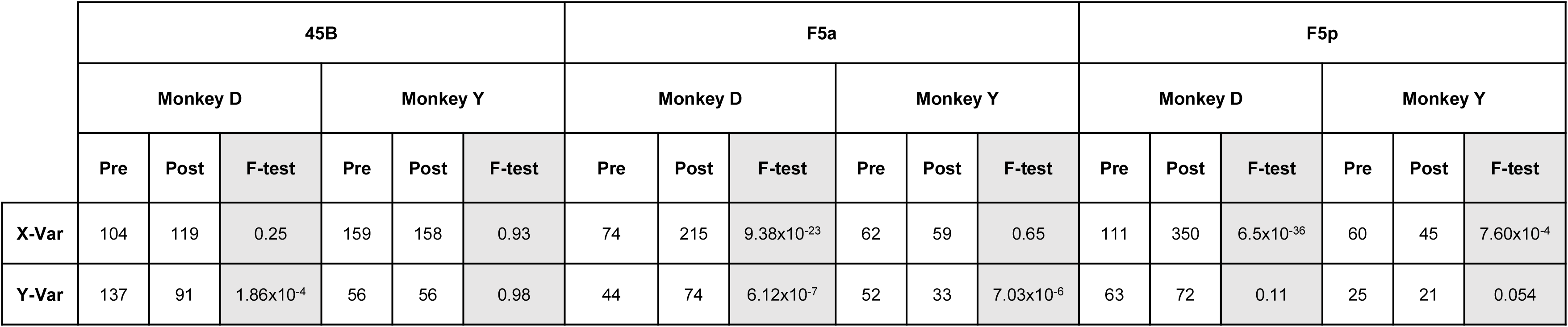
Eye position variance in areas 45B, F5a and F5p, before and after muscimol injection. Pre and Post columns represents for each monkey and each areas the variance in the position of one eye (X- and Y-coordinates). F-test column represents instead the p-value obtained with F-statistic comparing the eye position variability Pre and Post muscimol.

## Discussion

We investigated the causal role of area 45B, F5a and F5p in the grasping network during a VGG task. The monkeys were trained to reach and grasp two spheres of different sizes presented by a robot, at a reachable distance. Reversible inactivation of all three frontal areas induced a significant effect on grasping, suggesting a causal role of each area in visually guided object grasping.

Our results are highly relevant for understanding the cortical network subserving object grasping. The deficit induced by F5p inactivation corroborates previous findings (Fogassi et al., 2001) and confirms the role of F5p as the primary source of commands for hand control exerted by M1. Much less is known about the anterior subsector of PMv, F5a. Guided by a previous monkey fMRI study (Joly et al., 2009), Theys et al. (2012a-b) reported robust selectivity for the 3D structure of surfaces defined by binocular disparity in F5a, similar to its input area AIP (Srivastava et al., 2009; Premereur et al., 2015). Moreover, most of these 3Dstructure selective neurons were also active during object grasping in the light, but not in the dark, which was the first demonstration of visual-dominant neurons in F5. Especially noteworthy in the context of the current study was the observation that visual-dominant F5a neurons were frequently localized in close proximity with visuomotor and motor-dominant neurons. Thus, the inactivation of F5a most likely impaired the visual and visuomotor inputs to F5p, which caused the grasping deficit (albeit rarely accompanied with a change in the preshaping of the hand). The deficit after area 45B inactivation – which was almost as large as the one after F5a or F5p inactivation – is more difficult to interpret. Since area 45B is not on the ‘direct’ visual-to-motor pathway aAIP – F5a – F5p – M1 and given its anatomical location anterior to the arcuate sulcus and neighboring the FEF, the most straightforward interpretation is that this area plays a role in oculomotor control during grasping. However, it should be noted that in our experiments, the animals did not have to make a saccadic eye movement towards the object since it appeared in central vision. In addition, 45B neurons responded to the object that was fixated and remained active throughout the trial in the absence of any eye movement. Finally, we previously observed (Caprara & Janssen, unpublished data) that 45B neurons are almost not active during object grasping in the dark. Therefore, the oculomotor role of 45B should not be interpreted as the planning of an eye movement to to-be-grasped objects or object parts (Caprara et al., 2018). Instead, our results may suggest that the elevated 45B activity during visually guided object grasping is important for processing the visual feedback related to the position of the object and the hand approaching the object. In this sense, area 45B could be important for eye-hand coordination during grasping, a process that is also crucial in humans since the gaze guides the hand towards appropriate grasping contact points (Johansson et al., 2001). Future experiments using reversible inactivation with high temporal resolution (e.g. optogenetics) may further clarify how area 45B contributes to object grasping under visual guidance.

The pattern of anatomical connectivity of area 45B is also relevant for understanding its role in object grasping (Gerbella et al., 2010). Besides many prefrontal connections (area 46 and the FEF), area 45B is connected to the anterior IPS (described as LIP in Gerbella et al., 2010 but most likely corresponding to pAIP) and to parts of the inferotemporal cortex primarily in the lower bank of the rostral Superior Temporal Sulcus (TEa/m). However, area 45B seemed to have very little connections with F5a or F5p (also described in Gerbella et al. 2011), suggesting that the grasping deficit we observed after 45B inactivation might originate through other prefrontal areas such as area 12. The latter area has also been implicated in grasping (Hoshi et al., 2000; Borra et al., 2011; Simone et al., 2015; Simone et al., 2017) and is connected to F5a.

A number of alternative explanations and potential confounds also have to be considered. When inactivating frontal cortical areas, an attentional effect is always possible. We believe it is highly unlikely that we induced an attentional deficit when injecting muscimol in area 45B. Firstly, only one object appeared in foveal vision, hence any competition between different potential target objects was absent. Secondly, an attention deficit would most likely induce an increase in reaction time (i.e. the time between the go-cue and the lift of the hand), whereas we consistently measured significant effects on the grasping time (i.e. the time between the lift of the hand and the pull of the object) in all three areas. In fact, our control injection in area 46 – a key area for selective attention (Baldauf and Desimone, 2014; Gregoriou et al., 2014) – indeed caused longer reaction times with a much milder effect on the grasping times (the significant effect on the total grasping time was caused mainly by the reaction time effect). Spillover of muscimol from the 45B site to other parts of ventral premotor cortex is also a genuine concern when comparing effects of inactivation of neighboring areas. First, we verified the center of inactivation with a recording microelectrode at that location during anatomical MRI. In addition, we imaged the spread of the injected volume using dotarem injections and MRI, which provide an accurate estimate of the muscimol spread (Wilke et al., 2012). Our injections were always located at the targeted sites, and although the MRI suggests some spread around the injection site in 45B, it is highly unlikely that muscimol could cross the sulcus and affected the posterior bank of the arcuate sulcus in either F5a or F5p. Even the overlap between the F5a and the F5p injections was minimal. Finally, our conclusions only pertain to the specific setup we used: grasping a single, stable object in foveal vision during visual guidance. This experimental task is undoubtedly more relevant for everyday behavior since we almost never grasp objects in the dark. In fact, we predict that the inactivation of area 45B will not induce a grasping deficit when grasping the object in complete darkness since no visual feedback from the hand approaching the object is present in those circumstances.

We did not test whether smaller volumes would also induce a measurable grasping deficit. The use of a standard volume of 4 μl was motivated by the observation that the frontal areas we targeted extend over 3-4 mm when mapped with single-cell recordings. Moreover, a previous study (Van Dromme et al., 2016) also showed consistent behavioral and functional (fMRI activations) effects with muscimol injections of 4 μl. We also did not track the position of all fingers during grasping. Despite the very small size of one of the two spheres (1 cm diameter), the monkeys naturally used a combination of pad-to-side/scissor grip type for both objects, adapting the thumb and the hand aperture on the object, according to its dimension. The monkeys’ choice of not using a precision grip could be due to the distance of the objects, which although reachable, required a more powerful grip. It is possible that with a larger subset of stimuli we could have observed stronger deficits, or larger differences in the pattern of impairments in the three areas we tested.

In conclusion, our results provide the first causal evidence that area 45B and F5a are involved in visually guided object grasping. Future experiments combining single-cell recordings, functional imaging and causal perturbation methods may further clarify the flow of information in the frontal grasping network.

## Notes

### Competing Interest Statement

The authors have declared no competing interest.

